# Interferon receptor-deficient mice are susceptible to eschar-associated rickettsiosis

**DOI:** 10.1101/2020.09.23.310409

**Authors:** Thomas P. Burke, Patrik Engström, Cuong J. Tran, Dustin R. Glasner, Diego A. Espinosa, Eva Harris, Matthew D. Welch

## Abstract

*Rickettsia* are arthropod-borne pathogens that cause severe human disease worldwide. The spotted fever group (SFG) pathogen *Rickettsia parkeri* elicits skin lesion (eschar) formation in humans after tick bite. However, intradermal inoculation of inbred mice with millions of bacteria fails to elicit eschar formation or disseminated disease, hindering investigations into understanding eschar-associated rickettsiosis. Here, we report that intradermal infection of mice deficient for both interferon receptors (*Ifnar*^*-/-*^*Ifngr*^*-/-*^) with *R. parkeri* causes eschar formation, recapitulating the hallmark clinical feature of human disease. Intradermal infection with doses that recapitulate tick infestation caused eschar formation and lethality, including with as few as 10 bacteria. Using this model, we found that the actin-based motility protein Sca2 is required for *R. parkeri* dissemination from the skin to internal organs and for causing lethal disease, and that the abundant *R. parkeri* outer membrane protein OmpB contributes to eschar formation. We also found that immunizing mice with *sca2* and *ompB* mutant *R. parkeri* protects against subsequent rechallenge with wild-type bacteria, revealing live-attenuated vaccine candidates. Thus, interferon receptor-deficient mice are a tractable model to investigate rickettsiosis, bacterial virulence factors, and immunity. Our results suggest that differences in interferon signaling in the skin between mice and humans may explain the discrepancy in susceptibility to SFG *Rickettsia*.

## Introduction

Obligate cytosolic bacterial pathogens in the family Rickettsiaceae are a diverse group of arthropod-borne microbes that cause severe human disease worldwide, including spotted fever, scrub typhus, and typhus^1–3^. Human disease caused by the tick-borne spotted fever group (SFG) pathogen *Rickettsia parkeri* is characterized by an eschar at the infection site, generalized rash, headache, fatigue, and fever^4^. There is no approved vaccine for *R. parkeri* or for the more virulent rickettsial pathogens that can cause fatal or latent disease^5^. Moreover, many critical aspects of disease caused by obligate cytosolic bacterial pathogens, including the mechanisms of virulence and immunity, remain unknown, as there are no SFG pathogens that can be handled under biosafety level 2 (BSL2) conditions with corresponding mouse models that recapitulate key features of human disease^5–7^.

*R. parkeri* is genetically similar to the more virulent human pathogens *R. rickettsii* and *R. conorii*^8,9^, and it can be handled under BSL2 conditions. Moreover, mutants can be generated using transposon mutagenesis^10,11^, and small rodents including mice are natural reservoirs for *R. parkeri*^12–15^. Thus, a mouse model for *R. parkeri* that recapitulates key features of human infection would greatly enhance investigations into understanding rickettsial disease. However, inbred mice including C57BL/6 and BALB/c develop no or minor skin lesions upon intradermal (i.d.) infection with millions of *R. parkeri*^6^. C3H/HEJ mice, which harbor a mutation in the gene encoding Toll-like receptor 4 (TLR4), the receptor for extracellular lipopolysaccharide (LPS), have been proposed as models for *R. parkeri*, yet they do not develop disseminated disease and only develop minor skin lesions upon i.d. inoculation^6^. C57BL/6 mice have also been proposed as models for *R. parkeri* upon intravenous (i.v.) delivery of 10^8^ bacteria^16^. However, this dose is substantially higher than the number of *R. parkeri* found in tick saliva or tick salivary glands^17^, and considerable effort is required to generate and concentrate this number of bacteria. An improved mouse model to investigate *R. parkeri* would greatly increase the ability to investigate virulence mechanisms, the host response to infection, and human rickettsial disease.

Towards better understanding the host response to *R. parkeri* infection, we recently investigated the relationship between *R. parkeri* and interferons (IFNs), which are ubiquitous signaling molecules of the innate immune system that mobilize the cytosol to an antimicrobial state. Type I IFN (IFN-I) generally restricts viral replication, whereas IFN-*γ* generally restricts intracellular bacterial pathogens^18–20^. We observed that mice lacking either gene encoding the receptors for IFN-I (*Ifnar*) or IFN-*γ* (*Ifngr*) are resistant to i.v. infection with *R. parkeri*, whereas double mutant *Ifnar*^*-/-*^*Ifngr*^*-/-*^ mice succumb^21^. This demonstrates that IFNs redundantly protect against systemic *R. parkeri*. However, the i.v. infection route does not recapitulate eschar formation or mimic the natural route of dissemination. Moreover, *Ifnar*^*-/-*^*Ifngr*^*-/-*^ mice are resistant to i.v. infection with 10^5^ bacteria, which may exceed the amount delivered upon tick infestation. Further investigations into whether IFNs redundantly protect against *R. parkeri* in the skin may improve the mouse model for SFG *Rickettsia*.

A robust mouse model would allow for more detailed investigations into rickettsial virulence factors. One virulence mechanism shared by divergent cytosolic bacterial pathogens including *Rickettsia, Listeria, Burkholderia, Mycobacterium*, and *Shigella* species, is the ability to undergo actin-based motility, which facilitates cell to cell spread^22,23^. However, the pathogenic role for many actin-based motility factors *in vivo* remains poorly understood. *R. parkeri* actin-based motility differs from that of other pathogens in that it occurs in two phases, one that requires the RickA protein^10,24^ and the other that requires the Sca2 protein^10,25^. Only Sca2 is required for efficient cell to cell spread, although it is not required for replication in epithelial cells or for avoiding antimicrobial autophagy^10,25–27^. *sca2* mutant *R. rickettsii* elicit reduced fever in guinea pigs as compared with wild-type (WT) *R. rickettsii*^25^, yet the explanation for reduced fever and the pathogenic role for Sca2 *in vivo* remains unclear. Additionally, Sca2 is not essential for dissemination of *R. parkeri* within ticks^28^. A second virulence strategy employed by intracellular pathogens is the ability to avoid autophagy, which for *R. parkeri* requires outer membrane protein B (OmpB)^27^. OmpB is important for *R. parkeri* colonization of internal organs in WT mice and for causing lethal disease in IFN receptor-deficient mice after i.v. infection^21,27^; however, the role for OmpB in *R. parkeri* pathogenesis remains unknown upon i.d. infection. Therefore, unresolved questions remain regarding how Sca2 and OmpB enhance rickettsial pathogenesis.

Here, we use IFN receptor-deficient mice to examine the effects of i.d. inoculation of *R. parkeri*, mimicking the natural route of infection. We observe skin lesions that appear similar to human eschars, as well as disseminated lethal disease with as few as 10 bacteria. Using this model, we find that Sca2 promotes dissemination and is required for causing lethality, and that OmpB contributes to eschar formation. Finally, we demonstrate that immunization with *sca2* or *ompB* mutant *R. parkeri* protects IFN receptor-deficient mice against subsequent challenge with WT bacteria, revealing live-attenuated vaccine candidates. Our study establishes a mouse model to investigate numerous aspects of *Rickettsia* pathogenesis, including eschar formation, virulence factors, and immunity. More broadly, this work also reveals that a potent, redundant IFN response protects mice from eschar-associated rickettsiosis.

## Results

### I.d. infection of *Ifnar*^*-/-*^*Ifngr*^*-/-*^ mice causes lethal disease and skin lesions that are grossly similar to human eschars

Although i.v. delivery can recapitulate an immediate systemic disease for many pathogens, it does not mimic the natural route of infection for tick-borne pathogens. In contrast, i.d. delivery better mimics the natural route of infection and allows for investigations into dissemination from the initial infection site to internal organs. We therefore sought to develop an i.d. murine infection model to better recapitulate the natural route of tick-borne *R. parkeri* infection. WT, *Tlr4*^*-/-*^, *Ifnar*^*-/-*^, *Ifngr*^*-/-*^, and *Ifnar*^*-/-*^*Ifngr*^*-/-*^ C57BL/6J mice, as well as outbred CD-1 mice, were infected i.d. with 10^7^ WT *R. parkeri* and monitored over time. No or minor dermal lesions appeared at the site of infection in WT, *Tlr4*^-/-^, *Ifnar*^*-/-*^, or *Ifngr*^-/-^ C57BL/6J mice or CD-1 mice (**Fig. 1a, Fig. S1a**). In contrast, double mutant *Ifnar*^*-/-*^*Ifngr*^*-/-*^ C57BL/6J mice developed large necrotic skin lesions (**Fig. 1b**) that appeared grossly similar to human eschars (**Fig. 1c**). In some cases, tails of *Ifnar*^*-/-*^*Ifngr*^*-/-*^ or *Ifngr*^*-/-*^ mutant mice became inflamed after i.d. or i.v. infection (**Fig. S1b**). These findings demonstrate that interferons redundantly control disease caused by *R. parkeri* in the skin and that i.d. infection of *Ifnar*^*-/-*^*Ifngr*^*-/-*^ mice recapitulates the hallmark manifestation of human disease caused by *R. parkeri*.

**Figure 1:**
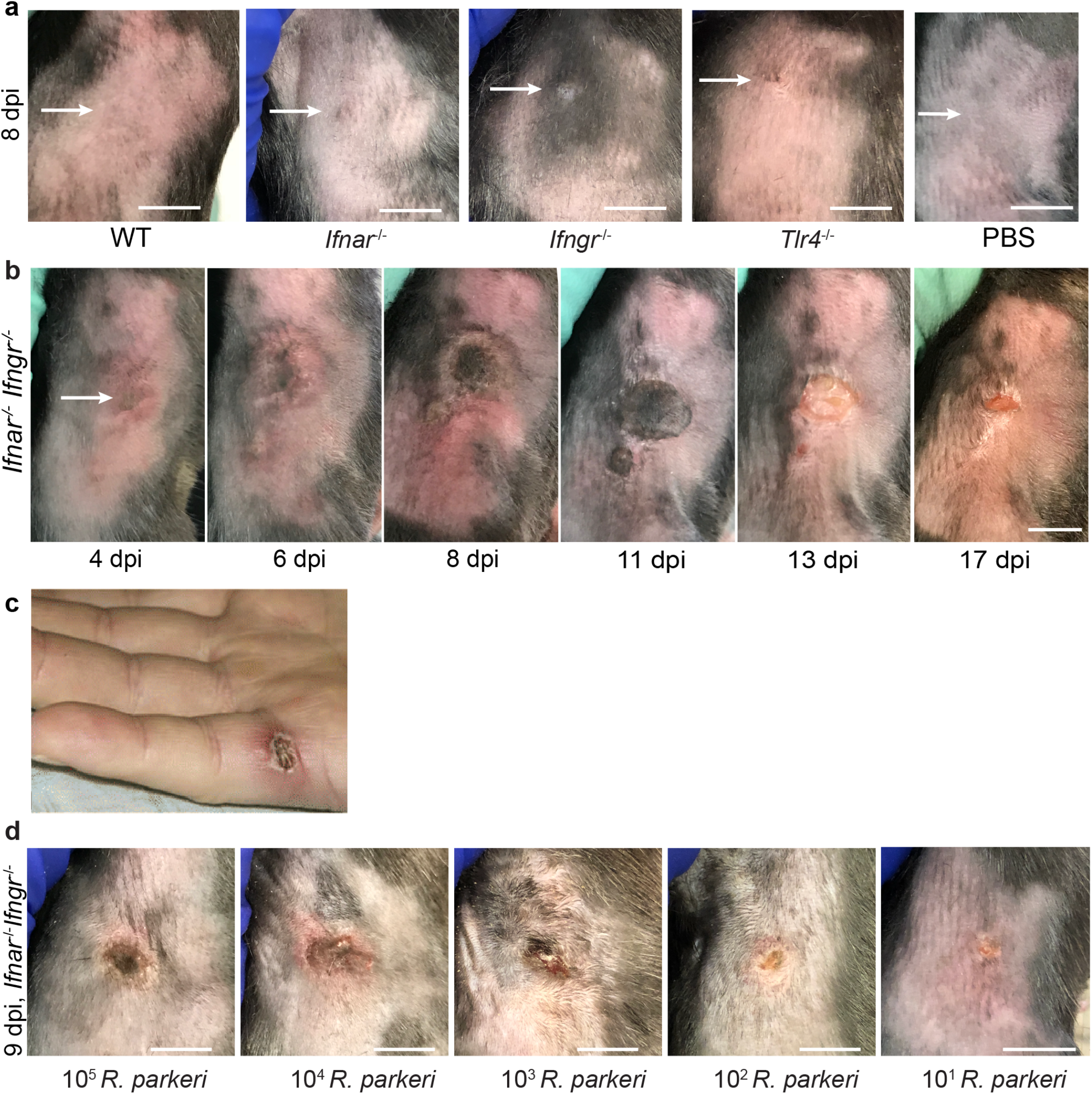
I.d. infection of *Ifnar*^*-/-*^*Ifngr*^*-/-*^ mice with *R. parkeri* elicits skin lesions that are grossly similar to human eschars. **a**) Representative images of WT, *Ifnar*^*-/-*^, *Ifngr*^*-/-*^, and *Tlr4*^*-/-*^ mice, infected intradermally with 10^7^ WT *R. parkeri* at 8 dpi and WT mice injected with PBS. White arrows indicate the infection site on the right flank of the mouse. Scale bar, 1 cm. Data are representative of three independent experiments. **b**) Representative images of an *Ifnar*^*-/-*^*Ifngr*^*-/-*^ mouse after i.d. inoculation with 10^7^ *R. parkeri*. Data are representative of 3 independent experiments. The white arrow indicates the injection site on the right flank of the mouse. Scale bar, 1 cm. **c**) Gross pathology of a human *R. parkeri* infection, from Paddock *et al*^4^. **d**) Representative images of *Ifnar*^*-/-*^*Ifngr*^*-/-*^ mice infected intradermally with the indicated amounts of WT *R. parkeri* at 9 d.p.i. Scale bar, 1 cm. Data are representative of two independent experiments.

Our previous observations using the i.v. route revealed dose-dependent lethality in *Ifnar*^*-/-*^*Ifngr*^*-/-*^ mice, with 10^7^ *R. parkeri* eliciting 100% lethality and 10^5^ *R. parkeri* eliciting no lethality^21^. *R. parkeri* are present in tick saliva at a concentration of approximately 10^4^ per 1 μl, and approximately 10^7^ *R. parkeri* are found in tick salivary glands^17^. However, the number of bacteria delivered from tick infestation likely varies depending on many factors, and we therefore sought to examine the effects of different doses of *R. parkeri* upon i.d. infection of *Ifnar*^*-/-*^*Ifngr*^*-/-*^ mice. We observed skin lesion formation at all infectious doses, from 10^7^ to 10 bacteria (**Fig. 1d**), suggesting that i.d. infection of *Ifnar*^*-/-*^*Ifngr*^*-/-*^ mice elicits lesions with doses similar to what is delivered by tick infestation.

We next sought to quantitatively evaluate the effects of i.d. infection by monitoring animal weight, body temperature, the degree of lesion formation, and lethality. Intradermally-infected *Ifnar*^*-/-*^*Ifngr*^*-/-*^ mice lost significant body weight (**Fig. 2a; Fig. S2a**) and body temperature (**Fig. 2b**; animals were euthanized when body temperature fell below 90° F / 32.2° C) as compared with WT mice, whereas infected *Tlr4*^*-/-*^, *Ifnar*^*-/-*^, *Ifngr*^*-/-*^ mice did not. To evaluate lesion severity, we scored lesions upon infection with different doses of *R. parkeri*. Whereas 10^7^ bacteria elicited similar responses as 10^5^, 10^4^, 10^3^, and 10^2^ bacteria (**Fig. 2c**), lesions were less severe when mice were infected with 10^1^ bacteria compared with 10^7^ bacteria. If mice survived, lesions healed over the course of approximately 15-40 days post infection (d.p.i.) at all doses (**Fig. S2b**).

**Figure 2:**
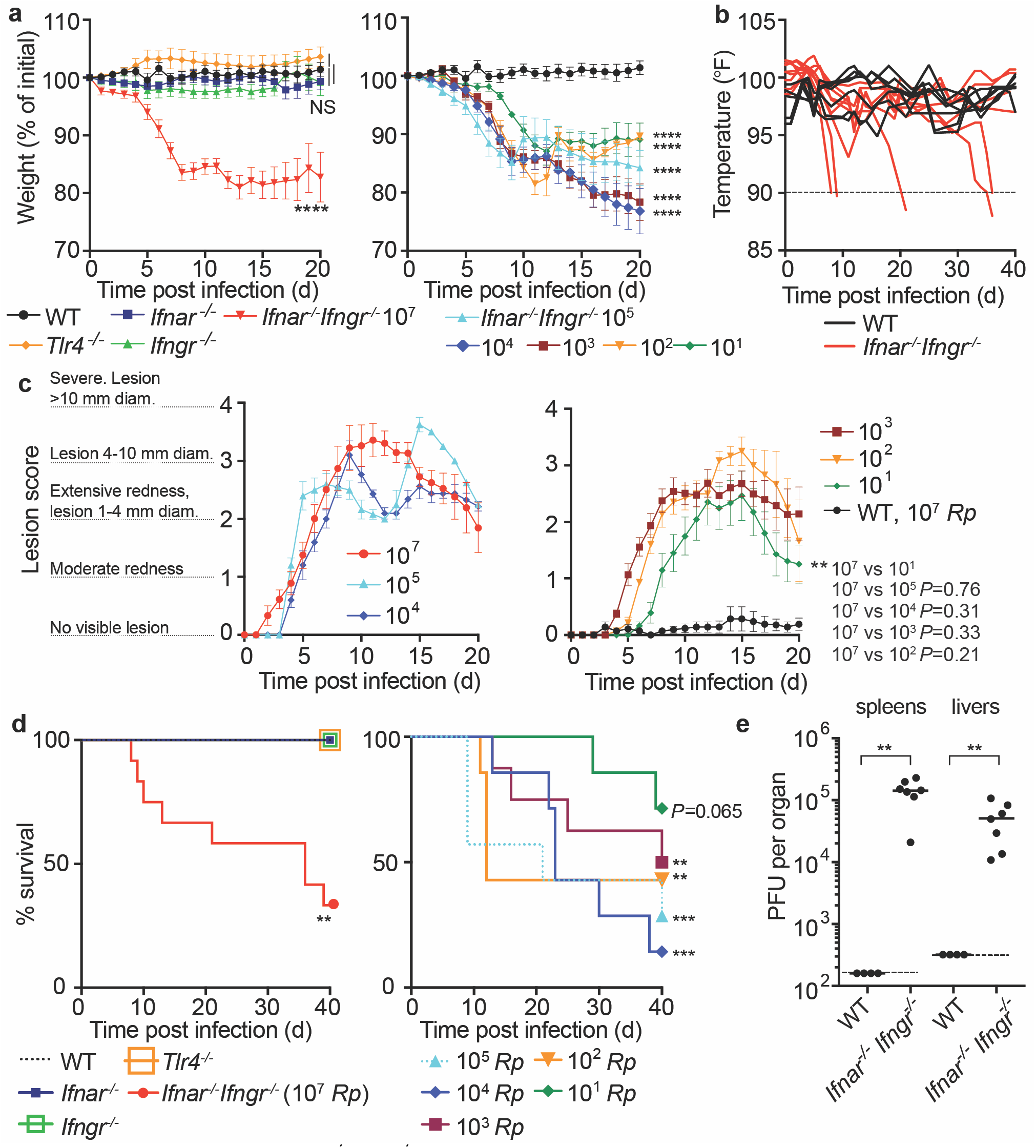
I.d. infection of *Ifnar*^*-/-*^*Ifngr*^*-/-*^ mice by *R. parkeri* elicits disseminated, lethal disease. **a**) Weight changes over time in mice infected i.d. with *R. parkeri*. Data are shown as a percentage change to initial weight. In the left panel, all mice were infected with 10^7^ *R. parkeri*; *n*=7 (WT), *n*=11 (*Ifnar*^*-/-*^), *n*=7 (*Ifngr*^*-/-*^), *n*=9 (*Ifnar*^-/-^*Ifngr*^-/-^) and 4 (*Tlr4*^-/-^) individual mice. In the right panel, *Ifnar*^-/-^*Ifngr*^-/-^ mice were infected with the indicated amounts of *R. parkeri*; *n*=7 (10^5^ *R. parkeri*), *n*=7 (10^4^ *R. parkeri*), *n*=8 (10^3^ *R. parkeri*), *n*=7 (10^2^ *R. parkeri*), *n*=7 (10^1^ *R. parkeri*) individual mice. WT data is the same in both panels. Data for each genotype are combined from two or three independent experiments. **b**) Temperature changes over time in mice intradermally infected with 10^7^ *R. parkeri*. Each line is an individual mouse. Mice were euthanized if their temperature fell below 90° F, as indicated by the dotted line. Data are the combination of three independent experiments with *n=*7 (WT) and 9 (*Ifnar*^*-/-*^*Ifngr*^*-/-*^) individual mice. **c**) Analysis of gross skin pathology after i.d. infection. *Ifnar*^*-/-*^*Ifngr*^*-/-*^ mice were infected with the indicated number of *R. parkeri* and monitored over time. WT mice were infected with 10^7^ *R. parkeri*. Data are the combination of three independent experiments for WT and the 10^7^ dose in *Ifnar*^*-/-*^ *Ifngr*^*-/-*^ mice; data for all other doses are the combination of two independent experiments. *n=*9 (10^7^), *n=*5 (10^5^), *n=*5 (10^4^), *n=*8 (10^3^), *n=*7 (10^2^), *n=*7 (10^1^), and *n=*7 (WT) individual mice. **d**) Mouse survival after i.d. infection with *R. parkeri*. In the left panel, all mice were infected with 10^7^ *R. parkeri*; *n*=7 (WT), *n*=11 (*Ifnar*^*-/-*^), *n*=7 (*Ifngr*^*-/-*^), *n*=4 *Tlr4*^*-/-*^, and *n*=12 (*Ifnar*^*-/-*^*Ifngr*^*-/-*^) individual mice. Data are the combination of three separate experiments for WT, *Ifnar*, and *Ifnar*^*-/-*^*Ifngr*^*-/-*^ and two separate experiments for *Ifngr*^*-/-*^ and *Tlr4*^*-/-*^. In the right panel, *Ifnar*^*-/-*^*Ifngr*^*-/-*^ mice were infected with the indicated amounts of *R. parkeri*. Data are the combination of two independent experiments; *n=*7 (10^5^), *n=*7 (10^4^), *n=*8 (10^3^), *n=*7 (10^2^), and *n=*7 (10^1^) individual mice. **e**) Bacterial burdens in organs of intradermally infected WT and *Ifnar*^*-/-*^ *Ifngr*^*-/-*^ mice. Mice were intradermally inoculated with 10^7^ *R. parkeri*, and spleens and livers were harvested and plated for p.f.u. at 72 h.p.i. Dotted lines indicate the limit of detection. Data are the combination of two independent experiments. *n*=4 (WT) and 7 (*Ifnar*^-/-^*Ifngr*^-/-^) individual mice. Data in **a, c** are the mean ± SEM. Statistical analyses in **a** used a two-way ANOVA where each group was compared to WT at t=20 d.p.i. Statistical analyses in **c** used a two-way ANOVA at t=20 d.p.i. Statistical analyses in **d** used a log-rank (Mantel-Cox) test to compare *Ifnar*^-/-^ to *Ifnar*^-/-^*Ifngr*^-/-^ at each dose. Statistical analysis in **e** used a two-tailed Mann-Whitney U test. NS, not significant; \*\**P*<0.01; \*\*\**P*<0.001; \*\*\*\**P*<0.0001.

To investigate whether i.d. infection by *R. parkeri* caused lethal disease, we monitored mouse survival over time. Upon i.d. delivery of 10^7^ *R. parkeri*, 8 of 12 *Ifnar*^*-/-*^*Ifngr*^*-/-*^ mice exhibited lethargy, paralysis, or body temperatures below 90° F, at which point they were euthanized, whereas delivery of the same dose of bacteria to WT and single mutant mice did not elicit lesions and all survived (**Fig. 2d**). Lower doses of *R. parkeri* also elicited body weight loss (**Fig. 2a**), body temperature loss (**Fig. S2c**), and lethal disease (**Fig. 2d**) in *Ifnar*^*-/-*^*Ifngr*^*-/-*^ mice. The cause of lethality in this model remains unclear and will require further investigation. Nevertheless, these findings reveal that i.d. infection can cause lethal disease in *Ifnar*^*-/-*^*Ifngr*^*-/-*^ mice with ∼10,000-fold lower dose of bacteria than i.v. infection.

It remained unclear whether i.d. infection could also be used to model dissemination from the skin to internal organs. We therefore evaluated bacterial burdens in spleens and livers of WT and *Ifnar*^*-/-*^*Ifngr*^*-/-*^ mice at 5 d.p.i. by measuring *R. parkeri* plaque-forming units (p.f.u.). Bacteria were not recoverable from spleens and livers of intradermally-infected WT mice, suggesting that they did not disseminate from the skin to internal organs in high numbers (**Fig. 2e**). In contrast, bacteria were recovered from spleens and livers of intradermally-infected *Ifnar*^*-/-*^*Ifngr*^*-/-*^ mice at 5 d.p.i. (**Fig. 2e**). This demonstrates that i.d. infection of *Ifnar*^*-/-*^*Ifngr*^*-/-*^ mice with *R. parkeri* causes systemic infection and can be used as a model for dissemination from the skin to internal organs.

### *Ifnar*^*-/-*^*Ifngr*^*-/-*^ mice do not succumb to intradermal infection with *sca2* mutant *R. parkeri*

Sca2 mediates actin-based motility in rickettsial pathogens; however, its contribution to virulence *in vivo* remains unclear. We examined if i.v. and i.d. infections of WT and *Ifnar*^*-/-*^*Ifngr*^*-/-*^ mice could reveal a pathogenic role for *R. parkeri* Sca2. Upon i.v. infection with 5 x 10^6^ bacteria (**Fig. 3a**) or 10^7^ bacteria (**Fig. 3b)**, we observed that *sca2*::Tn mutant *R. parkeri* caused reduced lethality compared to WT bacteria. Similarly, i.d. infection with *sca2*::Tn mutant bacteria elicited significantly less lethality (**Fig. 3c**) and weight loss (**Fig. 3d**) as compared to WT bacteria and no severe temperature loss (**Fig. S3a**). Although we sought to evaluate infection using a *sca2* complement strain of *R. parkeri*, our attempts to generate such a strain were unsuccessful. As an alternative strategy, we examined whether the transposon insertion itself had an effect on *R. parkeri* survival *in vivo*. We evaluated infection of an *R. parkeri* strain that harbors a transposon insertion in *MC1_RS08740* (previously annotated as *MC1_05535*), which has no known role in virulence^27^. I.v. infection with *MC1_RS08740*::Tn *R. parkeri* caused lethality to a similar degree as WT *R. parkeri* (**Fig. 3a**), demonstrating that the transposon likely does not significantly impact *R. parkeri* fitness *in vivo*. Together, these findings suggest that the actin-based motility factor Sca2 is required for causing lethal disease in *Ifnar*^*-/-*^*Ifngr*^*-/-*^ mice.

**Figure 3:**
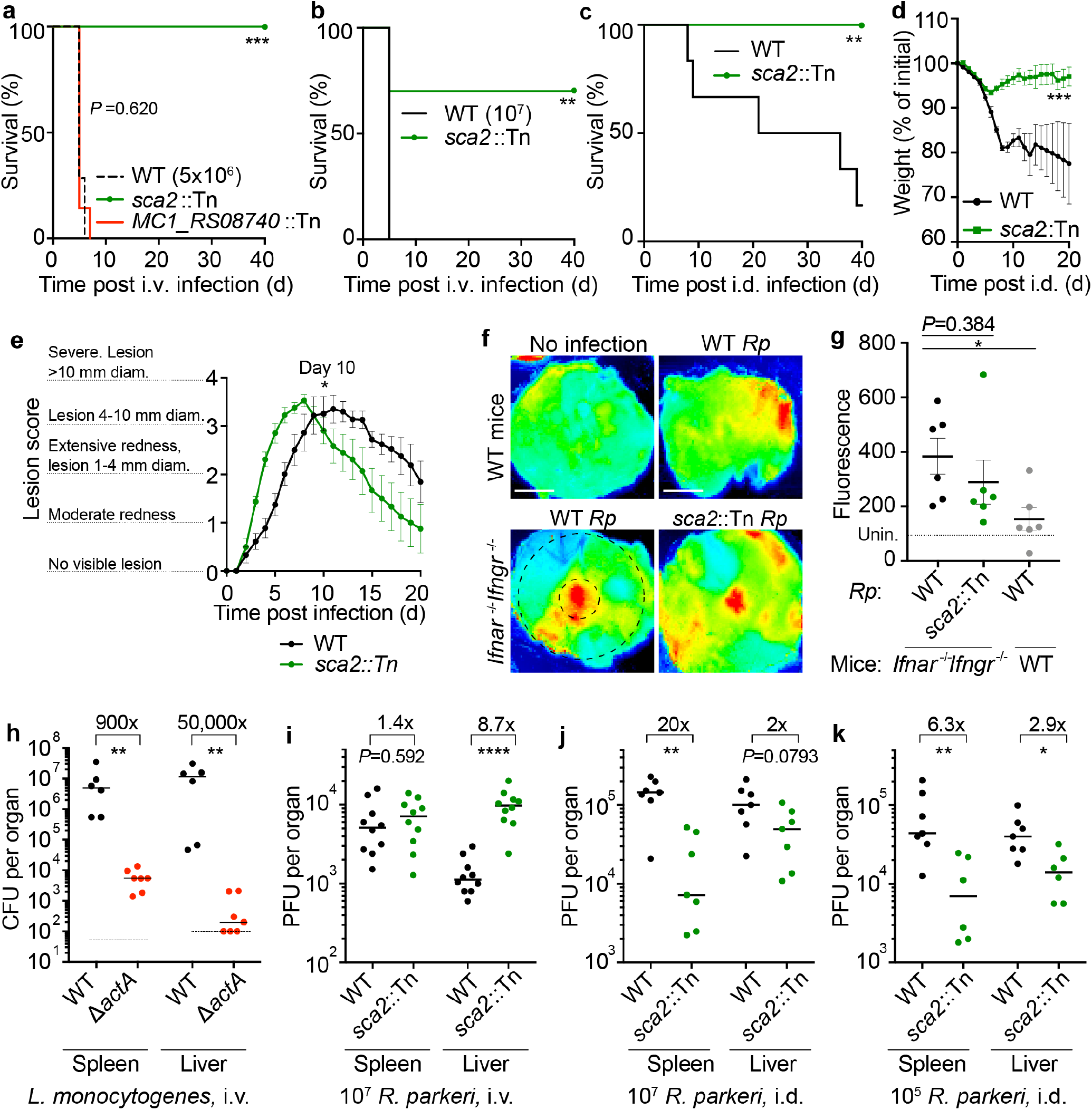
*R. parkeri* Sca2 contributes to dissemination from skin to spleens and livers. **a**) Survival of *Ifnar*^*-/-*^*Ifngr*^*-/-*^ mice upon i.v. infection with 5 x 10^6^ *R. parkeri. n*=7 (WT), 10 (*sca2*::Tn), and 7 (*MC1_RS08740*::Tn *R. parkeri*) individual mice. Data are the combination of two independent experiments. **b**) Survival of *Ifnar*^*-/-*^*Ifngr*^*-/-*^ mice upon i.v. infection with 10^7^ *R. parkeri. n*=7 (WT) and 10 (*sca2*::Tn) individual mice. Data are the combination of two independent experiments. **c**) Survival of *Ifnar*^*-/-*^*Ifngr*^*-/-*^ mice upon i.d. infection with 10^7^ *R. parkeri. n*=6 (WT) and 8 (*sca2*::Tn) individual mice. Data are the combination of two independent experiments. **d**) Weight changes of *Ifnar*^*-/-*^*Ifngr*^*-/-*^ mice upon i.d. infection with 10^7^ *R. parkeri. n*=6 (WT) and 8 (*sca2*::Tn) individual mice. Data are the combination of two independent experiments. **e**) Analysis of gross skin pathology after i.d. infection. *Ifnar*^*-/-*^*Ifngr*^*-/-*^ mice were infected with 10^7^ of the indicated strains of *R. parkeri* and monitored over time. *n*=9 (WT) and 8 (*sca2*::Tn) individual mice. Data are the combination of two independent experiments. **f**) Representative images of fluorescence in mouse skin after i.d. infection with 10^6^ *R. parkeri* and delivery of a fluorescent dextran, at 5 d.p.i. Scale bars, 1 cm. The larger black dashed circle represents the area that was measured for fluorescence for each sample, as indicated in **Fig. 3g** (∼80,000 pixels). The smaller black-dashed circle represents of the injection site area that was measured for fluorescence for each sample, as indicated in **Fig. S3** (∼7,800 pixels). **g**) Quantification of fluorescence in mouse skin after i.d. infection. Mice were infected with 10^7^ *R. parkeri*, and 150 µl fluorescent dextran was intravenously delivered at 5 d.p.i. Skin was harvested 2 h later, and fluorescence was measured using a fluorescence imager. Data indicate measurements of larger areas of skin, as indicated in **f** by the larger black circle. *n*=6 (WT *R. parkeri*) and *n*=6 (*sca2*::Tn *R. parkeri*) individual *Ifnar*^*-/-*^*Ifngr*^*-/-*^ mice; *n*=6 (WT *R. parkeri*) individual WT mice. For each experiment, the average of uninfected samples was normalized to 100; each sample was divided by the average for uninfected mice and multiplied by 100; the dotted horizontal line indicates 100 arbitrary units, corresponding to uninfected (unin.) mice. Data are the combination of two independent experiments. **h**) Quantification of *L. monocytogenes* abundance in organs of WT C57BL/6J mice upon i.v. infection with 10^4^ bacteria, at 72 h.p.i. Data are the combination of two independent experiments. n=6 (WT), n=7 (Δ*actA*) individual mice. **i**) Quantification of *R. parkeri* abundance in spleens and livers of WT C57BL/6J mice upon i.v. infection, at 72 h.p.i. Data are the combination of two independent experiments. *n*=10 (WT) and 10 (*Sca2*::Tn) individual mice. **j**) Quantification of *R. parkeri* abundance in organs upon i.d. infection with 10^7^ *R. parkeri. n*=7 (WT) and 7 (*sca2*::Tn) individual mice. Data are the combination of two independent experiments. Data for WT *R. parkeri* in *Ifnar*^*-/-*^*Ifngr*^*-/-*^ mice are the same as in **Fig. 2e. k**) Quantification of *R. parkeri* abundance in organs upon i.d. infection with 10^5^ *R. parkeri. n*=7 (WT) and 6 (*sca2*::Tn). Data are the combination of two independent experiments. Solid horizontal bars in **g** indicate means; solid horizontal bars in **h**-**k** indicate medians; error bars indicate SEM. Statistical analyses for survival in **a, b, c** used a log-rank (Mantel-Cox) test. Statistical analysis in **d** used a two-way ANOVA at t=20. Statistical analysis in **e** used a two-way ANOVA from 0 to 10 d.p.i. Statistical analyses in **g** used a two-tailed Student’s T test. Statistical analyses in **h, i, j, k** used a two-tailed Mann-Whitney U test. The fold change in **h, i, j, k** indicates differences of medians. \**P*<0.05; \*\**P*<0.01; \*\*\**P*<0.001; \*\*\*\**P*<0.0001.

### *Ifnar*^*-/-*^*Ifngr*^*-/-*^ mice exhibit similar skin lesion formation and vascular damage upon i.d. infection with WT and *sca2*::Tn *R. parkeri*

We next examined whether Sca2 facilitates *R. parkeri* dissemination throughout the skin and whether Sca2 is required for lesion formation. Unexpectedly, upon i.d. inoculation, *Ifnar*^*-/-*^*Ifngr*^*--*^ mice infected with *sca2*::Tn mutant bacteria developed skin lesions that were of similar severity to lesions caused by WT *R. parkeri*; however, the lesions elicited by *sca2* mutant bacteria appeared significantly earlier than lesions caused by WT bacteria (**Fig. 3e**). Further examinations will be required to better evaluate this observation; however, it may suggest that actin-based motility enables *R. parkeri* to avoid a rapid onset of inflammation in the skin. To evaluate *R. parkeri* dissemination within the skin, we used a fluorescence-based assay that measures vascular damage as a proxy for pathogen dissemination^29^. Mice were intradermally infected with WT and *sca2*::Tn *R. parkeri*. At 5 d.p.i., fluorescent dextran was intravenously delivered, and fluorescence was measured at the infection site (**Fig. 3f**, representative small black circle) and in the surrounding area (**Fig. 3f**, representative large black circle). No significant differences were observed when comparing WT and *sca2*::Tn *R. parkeri* infections in *Ifnar*^*-/-*^*Ifngr*^*-/-*^ mice using an infectious dose of 10^7^ *R. parkeri* in the larger surrounding area (**Fig. 3g**) or at the site of infection (**Fig. S4a**). Similar results were observed upon infection with 10^6^ or 10^5^ bacteria (**Fig. S4b**,**c**). However, significantly more fluorescence was observed in the skin of infected *Ifnar*^*-/-*^*Ifngr*^*-/-*^ mice as compared to WT mice (**Fig. 3g**), demonstrating that interferons protect against increased vascular permeability during *R. parkeri* infection. Together, the gross pathological analysis and fluorescence-based assay suggest that Sca2 likely does not significantly enhance *R. parkeri* dissemination in the skin during i.d. infection of *Ifnar*^*-/-*^*Ifngr*^*-/-*^ mice.

### *R. parkeri* Sca2 promotes dissemination from the skin to spleens and livers

Among the factors that mediate actin-based motility, the *L. monocytogenes* actin-based motility factor ActA is one of the best understood. ActA enables *L. monocytogenes* to spread from cell to cell^22,23^, escape antimicrobial autophagy^30–33^, proliferate in mouse organs after i.v. infection^34,35^, and cause lethal disease in mice^36,37^. We initially hypothesized that *R. parkeri* Sca2 plays a similar pathogenic role *in vivo* to ActA, which we found is required for bacterial survival in spleens and livers upon i.v. delivery (**Fig. 3h**), in agreement with previous experiments^34,35^. However, when we examined bacterial burdens upon i.v. infection of *Ifnar*^*-/-*^*Ifngr*^*-/-*^ mice with *R. parkeri*, similar amounts of WT and *sca2*::Tn bacteria were recovered in spleens (**Fig. 3i**). We were also surprised to find that significantly more *sca2*::Tn than WT *R. parkeri* were recovered in livers (**Fig. 3i**). The explanation for higher *sca2*::Tn burdens in livers remains unclear. Nevertheless, these data reveal that Sca2 is likely not essential for *R. parkeri* survival in blood, invasion of host cells, or intracellular survival in spleens and livers.

We next evaluated the role for Sca2 in *R. parkeri* dissemination by measuring p.f.u. in spleens and livers following i.d. infection of *Ifnar*^*-/-*^*Ifngr*^*-/-*^ mice. After i.d. infection, *sca2*::Tn mutant bacteria were ∼20-fold reduced in their abundance in spleens and ∼2-fold reduced in their abundance in livers as compared to WT *R. parkeri* (**Fig. 3j**). Similar results were seen upon i.d. infection with lower doses of *sca2*::Tn and WT bacteria (**Fig. 3k**). Together, these results suggest that Sca2 is required for *R. parkeri* dissemination from the skin to internal organs.

### *R. parkeri* actin-based motility does not contribute to avoiding innate immunity *in vitro*

Sca2-mediated actin-based motility is required for efficient plaque formation and cell to cell spread by *R. parkeri in vitro*^10,25^. However, it remains unclear if Sca2 enables *R. parkeri* to escape detection or restriction by innate immunity. The actin-based motility factor ActA enables *L. monocytogenes* to avoid autophagy^32,33^, and the antimicrobial guanylate binding proteins (GBPs) inhibit *Shigella flexneri* actin-based motility^38^. We therefore sought to evaluate whether Sca2-mediated actin-based motility enables *R. parkeri* to evade innate immunity *in vitro*. We found that the *sca2*::Tn mutant grew similarly to WT bacteria in endothelial cells (**Fig. S5a**), consistent with previous reports in epithelial cells^10,25^. We also examined whether Sca2 contributed to *R. parkeri* survival or growth in bone marrow-derived macrophages (BMDMs), which can restrict other *R. parkeri* mutants that grow normally in endothelial cells^27^. However, no significant difference in bacterial survival was observed between WT and *sca2*::Tn bacteria in BMDMs in the presence or absence of IFN-*β* (**Fig. S5b**). WT and *sca2* mutant *R. parkeri* also elicited similar amounts of host cell death (**Fig. S5c**) and IFN-I production (**Fig. S5d**). Moreover, we found that the anti-rickettsial factor GBP2 localized to the surface of *sca2*::Tn mutant *R. parkeri* at similar frequency as with WT bacteria in the presence or absence of IFN-*β* (**Fig. S5e,f**). Together, these data suggest that Sca2 does not significantly enhance the ability of *R. parkeri* to evade innate immunity *in vitro*.

### *Ifnar*^-/-^*Ifngr*^-/-^ mice exhibit less severe skin lesions upon infection with a highly attenuated *R. parkeri* mutant

Because *sca2* mutant *R. parkeri* showed no defect in eschar formation compared to WT, it remained unclear whether skin lesion formation in *Ifnar*^*-/-*^*Ifngr*^*-/-*^ mice was influenced by bacterial virulence factors. We therefore investigated i.d. infection with *ompB*::Tn^STOP^ *R. parkeri*, which harbors both a transposon and a stop codon in *ompB*^27^, and is severely attenuated *in vivo*^21,27^. In contrast with WT bacteria, i.d. infection of *Ifnar*^*-/-*^*Ifngr*^*-/-*^ mice with *ompB*::Tn^STOP^ *R. parkeri* caused no lethality (**Fig. 4a**) or reduced weight loss (**Fig. 4b**). The *ompB*::Tn^STOP^ mutant *R. parkeri* also caused significantly less severe skin lesions than WT bacteria (**Fig. 4c**). These findings suggest that *Ifnar*^*-/-*^*Ifngr*^*-/-*^ mice can be used as a model to identify bacterial genes important for eschar formation.

**Figure 4:**
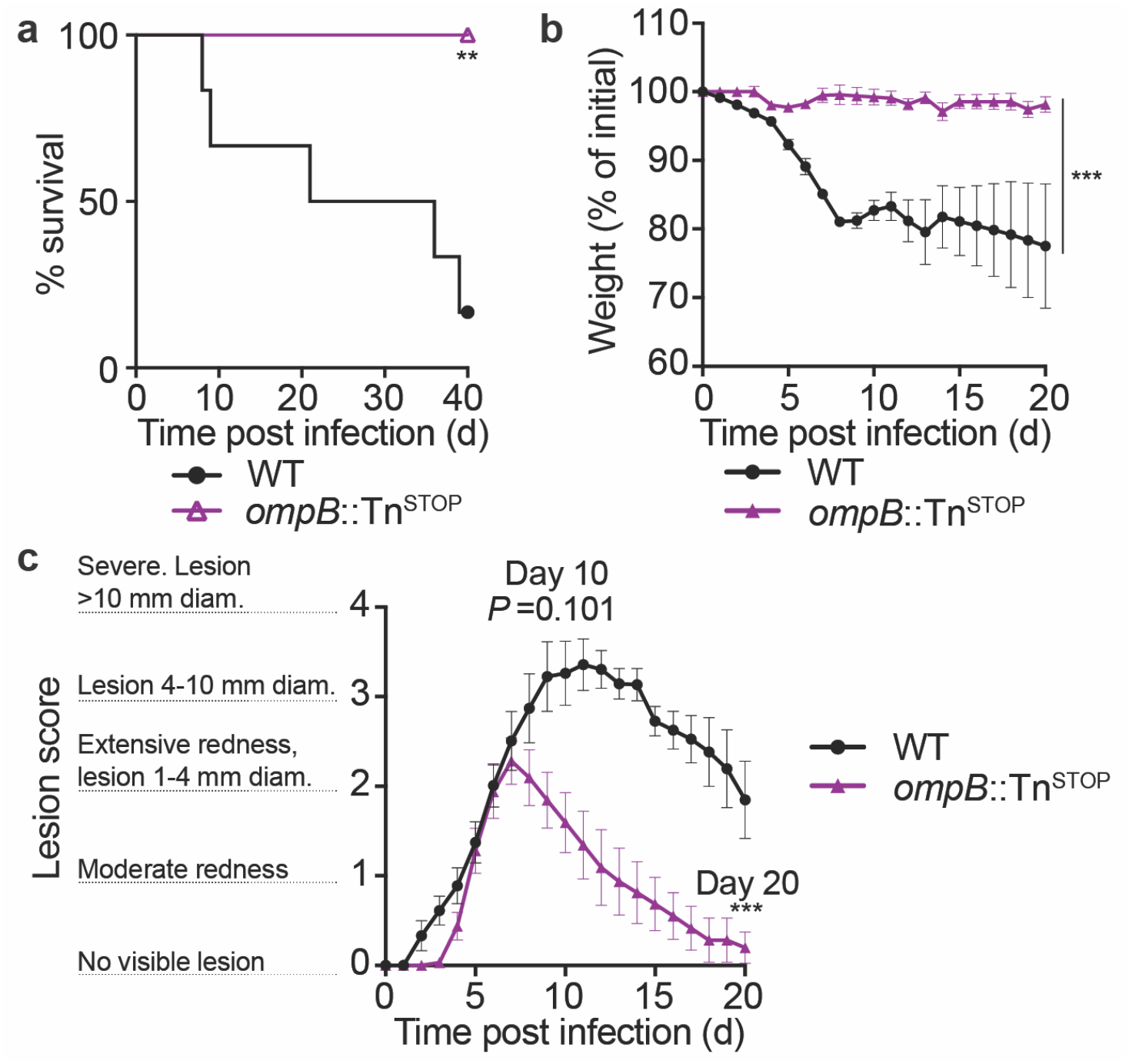
*ompB* mutant *R. parkeri* elicit no lethality and reduced skin lesion formation in *Ifnar*^-/-^ *Ifngr*^-/-^ mice. **a**) Survival of *Ifnar*^*-/-*^*Ifngr*^*-/-*^ mice upon i.d. infection with 10^7^ *R. parkeri. n*=6 (WT) and 8 (*ompB*::Tn^STOP^) individual mice. Data are the combination of two independent experiments. Data for WT are the same as in Fig. 3c. **b**) Weight changes of *Ifnar*^*-/-*^*Ifngr*^*-/-*^ mice upon i.d. infection with 10^7^ *R. parkeri. n*=6 (WT) and 8 (*ompB*::Tn^STOP^) individual mice. Data are the combination of two independent experiments. Data for WT are the same as in Fig. 3d. **c**) Analysis of gross skin pathology after i.d. infection. *Ifnar*^*-/-*^*Ifngr*^*-/-*^ mice were infected with 10^7^ of the indicated strains of *R. parkeri* and monitored over time. *n*=9 (WT) and 8 (*ompB*::Tn^STOP^) individual mice. Data are the combination of two independent experiments. Data for WT are the same as in Fig. 3e. Error bars indicate SEM. Statistical analyses in **a** used a log-rank (Mantel-Cox) test. Statistical analysis in **b** used a two-way ANOVA from 0 to 20 d.p.i. Statistical analysis in **c** used a two-way ANOVA from 0 to 10 and 20 d.p.i. \*\**P*<0.01; \*\*\**P*<0.001.

### Immunizing *Ifnar*^*-/-*^*Ifngr*^*-/-*^ mice with attenuated *R. parkeri* mutants protects against subsequent rechallenge

There is currently no available vaccine to protect against SFG *Rickettsia*, which can cause severe and lethal human disease^5,39^, and investigations into identifying live attenuated vaccine candidates has been hindered by the lack of robust animal models. We therefore examined whether immunization with attenuated *R. parkeri* mutants would protect against subsequent re-challenge with a lethal dose of WT bacteria. *Ifnar*^*-/-*^*Ifngr*^*-/-*^ mice were immunized i.v. with 5 x 10^6^ *sca2*::Tn or *ompB*::Tn^STOP^ *R. parkeri* and 40 d later were intravenously re-challenged with 10^7^ WT *R. parkeri*, which is approximately 10-times a 50% lethal dose (LD_50_)^21^. All mice immunized with *sca2* or *ompB* mutant *R. parkeri* survived, whereas all naïve mice succumbed by 6 d.p.i. (**Fig. 5a**). Upon rechallenge, mice immunized with *ompB* and *sca2* mutants also did not lose significant weight (**Fig. 5b**) or body temperature (**Fig. 5c**). These data indicate that attenuated *R. parkeri* mutants elicit a robust protective immune response, and that *Ifnar*^*-/-*^*Ifngr*^*-/-*^ mice may serve as tools to develop live attenuated *R. parkeri* vaccine candidates.

**Figure 5:**
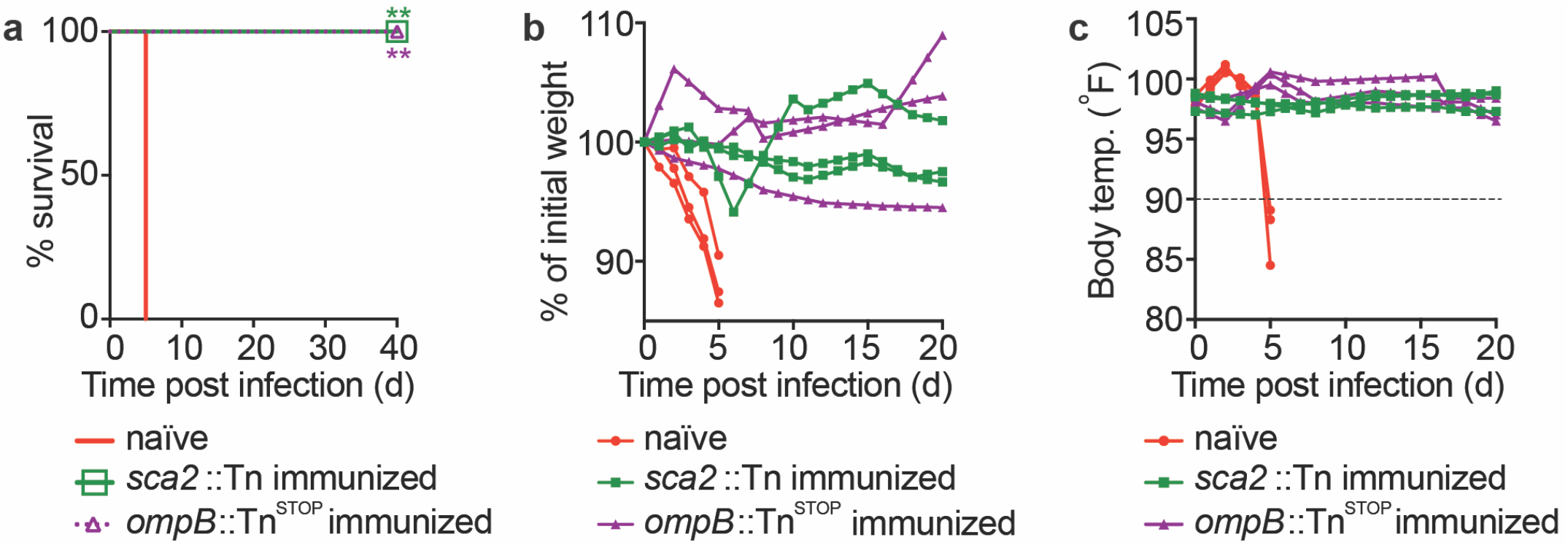
*ompB* and *sca2* mutant *R. parkeri* elicit immunity in *Ifnar*^*-/-*^*Ifngr*^*-/-*^ mice. **a**) Survival of immunized and naïve *Ifnar*^*-/-*^*Ifngr*^*-/-*^ mice upon i.v. *R. parkeri* infection. Immunized mice were first infected with 5×10^6^ *sca2*::Tn or 10^7^ *ompB*:Tn^STOP^ *R. parkeri* and were re-challenged 40 d later with 10^7^ WT *R. parkeri. n*=6 (naïve); *n*=5 (*sca2*::Tn immunized); *n*=5 (*ompB*::Tn^STOP^ immunized) individual mice. Data are the combination of two independent experiments. **b**) Weight changes over time in mice infected i.d. with 10^7^ *R. parkeri*. Data are representative of two independent experiments. *n*=3 (naïve); *n*=3 (*sca2*::Tn immunized); *n*=3 (*ompB*::Tn^stop^ immunized) individual mice. Each line represents an individual mouse. **c**) Temperature changes over time in mice infected i.d. with 10^7^ *R. parkeri*. Data are representative from two independent experiments. *n*=3 (naïve); *n*=3 (*sca2*::Tn immunized); *n*=3 (*ompB*::Tn^STOP^ immunized) individual mice. Each line represents an individual mouse. Statistical analyses in **a** used a log-rank (Mantel-Cox) test to compare each group of immunized mice to naïve mice. \*\**P*<0.01.

## Discussion

In this study, we show that IFN-I and IFN-*γ* redundantly protect inbred mice from eschar-associated rickettsiosis and disseminated disease by *R. parkeri*. Eschar formation is the hallmark clinical feature of human disease caused by *R. parkeri*^4^, and thus these findings suggest that the striking difference between human and mouse susceptibilities to *R. parkeri* may be due to IFN signaling in the skin. Using this mouse model, we uncover a role for *R. parkeri* Sca2 in dissemination, for OmpB in skin lesion formation, and for both proteins in causing lethal disease. We further demonstrate that attenuated *R. parkeri* mutants elicit long-lasting immunity, revealing live attenuated vaccine candidates. Obligate cytosolic bacterial pathogens cause a variety of severe human diseases on six continents^1,40^, and the animal model described here will facilitate future investigations into *R. parkeri* virulence factors, the host response to infection, the molecular determinants of human disease, and propagation of tick-borne pathogens in wildlife reservoirs.

Our finding that i.d. infection of *Ifnar*^*-/-*^*Ifngr*^*-/-*^ mice causes eschar formation, the hallmark of *R. parkeri* infection in humans^4^, may indicate that the human IFN response is less well adapted to control *R. parkeri* than in mice. Future investigations into the IFN-stimulated genes that restrict *R. parkeri* in mouse versus human cells may improve our understanding of human susceptibility to SFG *Rickettsia*. Additionally, our findings that OmpB promotes eschar formation demonstrates that *Ifnar*^*-/-*^*Ifngr*^*-/-*^ mice can be used to identify bacterial factors that are important for human disease manifestations. More broadly, investigating the IFN response in the skin may lead us to better understand diseases caused by other arthropod-borne pathogens. One example may be *Orientia tsutsugamushi*, the causative agent of scrub typhus^41^, a prevalent but poorly understood tropical disease endemic to Southeast Asia^1,42,43^. *O. tsutsugamushi* also causes eschar formation in humans, but inbred mice do not recapitulate eschar formation during *O. tsutsugamushi* infection^7^, similar to *R. parkeri*. A second example may be *Borrelia burgdorferi*, a tick-borne pathogen that causes a skin rash at the site of tick bite as a hallmark feature of Lyme disease^44^, the most prevalent tick-borne disease in the United States^44,45^. Existing mouse models also do not recapitulate skin rash formation following *B. burgdorferi* infection^46,47^. Further investigations into how IFNs protect the skin in mice may therefore reveal aspects of human disease caused by other arthropod-borne pathogens.

Our study further highlights the utility of mouse models that mimic natural routes of infection. Infection via the i.v. and intraperitoneal (i.p.) routes can mimic systemic disease, yet these are unnatural routes for many microbes, including food-borne, arthropod-borne, or aerosol-borne pathogens. Our observation that i.d. infection can cause lethal disease with as few as 10 bacteria, ∼10,000 fewer bacteria than i.v. infection^21^, suggests that *R. parkeri* may be highly adapted to reside in the skin. However, this model could be further improved by investigating the role for tick vector components in pathogenesis. Saliva from ticks, mosquitos, and sand flies enhances pathogenesis of arthropod-borne bacterial, viral, and parasitic pathogens^48–50^, and non-human primates inoculated with *R. parkeri* exhibit altered inflammatory responses when administered after tick-bite^51^. This may suggest a potential role for tick vector components such as tick saliva in *R. parkeri* pathogenesis. Developing improved murine infection models that mimic the natural route of infection, including with tick saliva or the tick vector, is critical to better understand the virulence and transmission of tick-borne pathogens.

Many *Rickettsia* species, as well as many facultative cytosolic pathogens including *L. monocytogenes*, undergo actin-based motility to spread from cell to cell. For *L. monocytogenes*, the actin-based motility factor ActA enables the pathogen to survive *in vivo*, as *actA* mutant bacteria are over 1,000-fold attenuated by measuring lethality^36,37^ and by enumerating bacteria in spleens and livers of mice after i.v. infection^34,35^. However, the pathogenic role for actin-based motility in the Rickettsiae has remained unclear. We find that Sca2 is not required for intracellular survival in organs upon i.v. infection of *Ifnar*^*-/-*^*Ifngr*^*-/-*^ mice, but rather, is required for dissemination from skin to internal organs and lethality upon i.d. infection. Consistent with an important role for Sca2 in pathogenesis, a previous study reported that i.v. infection of guinea pigs with *sca2* mutant *R. rickettsii* did not elicit fever^25^. Our results suggest that Sca2-mediated actin-based motility by *Rickettsia* may facilitate dissemination in host reservoirs, although we cannot rule out other roles for Sca2 that do not involve actin assembly. *R. prowazekii* and *R. typhi*, which cause severe human disease, encode a fragmented *sca2* gene^52^, and undergo no or dramatically reduced frequency of actin-based motility, respectively^53,54^. Although it remains unclear why some *Rickettsia* species lost the ability to undergo actin-based motility, Sca2 is dispensable for *R. parkeri* dissemination in the tick vector^28^, suggesting that actin-based motility may play a specific role in dissemination within mammalian hosts.

We find that *sca2* or *ompB* mutant *R. parkeri* elicit a robust protective immune response in *Ifnar*^*-/-*^*Ifngr*^*-/-*^ mice. These findings complement previous observations that *sca2* mutant *R. rickettsii* elicits antibody responses in guinea pigs^25^, and expands upon these findings by demonstrating protection from rechallenge and revealing additional vaccine candidates. There are currently limited vaccine candidates that protect against rickettsial disease^5^. Identifying new vaccine candidates may reveal avenues to protect against tick-borne infections and aerosolized *Rickettsia*, which are extremely virulent and potential bioterrorism agents^55^, as well as against Brill-Zinsser disease, caused by latent *R. prowazekii*^5^. Future studies exploring whether attenuated *R. parkeri* mutants provide immunity against other *Rickettsia* species are warranted to better define the mechanisms of protection. These findings on immunity may also help develop *R. parkeri* as an antigen delivery platform. *R. parkeri* resides directly in the host cytosol for days and could potentially be engineered to secrete foreign antigens for presentation by major histocompatibility complex I. In summary, the mouse model described here will facilitate future investigations into numerous aspects of *R. parkeri* infection, including actin-based motility and immunity, and may serve as model for other arthropod-borne pathogens.

## Methods

### Bacterial preparations

*R. parkeri* strain Portsmouth was originally obtained from Dr. Christopher Paddock (Centers for Disease Control and Prevention). To amplify *R. parkeri*, confluent monolayers of female African green monkey kidney epithelial Vero cells (obtained from UC Berkeley Cell Culture Facility, tested for mycoplasma contamination, and authenticated by mass spectrometry) were infected with 5 x 10^6^ *R. parkeri* per T175 flask. Vero cells were grown in DMEM (Gibco 11965-092) containing 4.5 gl^-1^ glucose and 2% fetal bovine serum (FBS; GemCell). Infected cells were scraped and collected at 5 or 6 d.p.i. when ∼90% of cells were rounded due to infection. Scraped cells were then centrifuged at 12,000*g* for 20 min at 4°C. Pelleted cells were resuspended in K-36 buffer (0.05 M KH_2_PO_4_, 0.05 M K_2_HPO_4_, 100 mM KCl, 15 mM NaCl, pH 7) and dounced for ∼40 strokes at 4°C. The suspension was then centrifuged at 200*g* for 5 min at 4°C to pellet host cell debris. Supernatant containing *R. parkeri* was overlaid on a 30% MD-76R (Merry X-Ray) gradient solution in ultracentrifuge tubes (Beckman/Coulter Cat 344058). Gradients were centrifuged at 18,000 r.p.m. in an SW-28 ultracentrifuge swinging bucket rotor (Beckman/Coulter) for 20 min at 4°C. These ‘30% prep’ bacterial pellets were resuspended in brain heart infusion (BHI) media (BD, 237500), aliquoted, and stored at -80°C. Titers were determined by plaque assays by serially diluting the *R. parkeri* in 6-well plates containing confluent Vero cells. Plates were spun for 5 min at 300*g* in an Eppendorf 5810R centrifuge and at 24 h post infection (h.p.i.); the media from each well was aspirated, and the wells were overlaid with 4 ml/well DMEM with 5% FBS and 0.7% agarose (Invitrogen, 16500-500). At 6 d.p.i., an overlay of 0.7% agarose in DMEM containing 2.5% neutral red (Sigma, N6264) was added. Plaques were then counted 24 h later. For infections with *ompB* mutant bacteria, the *ompB*^*STOP*^::Tn mutant was used, which contains a transposon and an upstream stop codon in *ompB*, as previously described^27^.

### Deriving bone marrow macrophages

For obtaining bone marrow, male or female mice were euthanized, and femurs, tibias, and fibulas were excised. Bones were sterilized with 70% ethanol and washed with BMDM media (20% FBS (HyClone), 0.1% β-mercaptoethanol, 1% sodium pyruvate, 10% conditioned supernatant from 3T3 fibroblasts, in DMEM (Gibco) with 4.5 gl^-1^ glucose and 100 μg/ml streptomycin and 100 U/ml penicillin), and ground with a mortar and pestle. Bone homogenate was passed through a 70 μm nylon cell strainer (Thermo Fisher Scientific, 08-771-2) for particulate removal. Filtrates were then centrifuged at 290*g* in an Eppendorf 5810R centrifuge for 8 min, supernatant was aspirated, and the pellet was resuspended in BMDM media. Cells were plated in 30 ml BMDM media in non-TC-treated 15 cm petri dishes at a ratio of 10 dishes per 2 femurs/tibias and incubated at 37° C. An additional 30 ml of BMDM media was added 3 d later. At 7 d the media was aspirated, 15 ml cold PBS (Gibco, 10010-023) was added, and cells were incubated at 4°C with for 10 min. BMDMs were scraped from the plate, collected in a 50 ml conical tube, and centrifuged at 290*g* for 5 min. PBS was aspirated, and cells were resuspended in BMDM media with 30% FBS and 10% DMSO at 10^7^ cells/ml. 1 ml aliquots were stored at -80° C for 24 h in Styrofoam boxes and then moved to long-term storage in liquid nitrogen.

### Infections *in vitro*

HMEC-1 cells (obtained from the UC Berkeley Cell Culture Facility and authenticated by short-tandem-repeat analysis) were passaged 2-3 times weekly and grown at 37° C with 5% CO_2_ in DMEM containing 10 mM L-glutamine (Sigma, M8537), supplemented with 10% heat-inactivated FBS (HyClone), 1 μg/mL hydrocortisone (Spectrum Chemical, CO137), 10 ng/mL epidermal growth factor (Thermo Fisher Scientific, CB40001; Corning cat. no. 354001), and 1.18 mg/mL sodium bicarbonate. HMEC media was prepared every 1-2 months, and aliquoted and stored at 4°C. To prepare HMEC-1 cells for infection, cells were treated with 0.25% trypsin-EDTA (Thermo Fisher Scientific); the number of cells was counted using a hemocytometer (Bright-Line), and 3 x 10^4^ cells were plated into 24-well plates 48 h prior to infection.

To plate macrophages for infection, BMDM aliquots were thawed on ice, diluted into 9 ml of DMEM, centrifuged at 290*g* for 5 min in an Eppendorf 5810R centrifuge, and the pellet was resuspended in 10 ml BMDM media without antibiotics. 5 x 10^5^ cells were plated into 24-well plates. Approximately 16 h later, “30% prep” *R. parkeri* were thawed on ice and diluted into fresh BMDM media to either 10^6^ p.f.u./ml or 2×10^5^ p.f.u./ml. Media was then aspirated from the BMDMs, replaced with 500 µl media containing *R. parkeri*, and plates were spun at 300*g* for 5 min in an Eppendorf 5810R centrifuge. Infected cells were incubated in a humidified CEDCO 1600 incubator set to 33°C and 5% CO_2_. Recombinant mouse IFN-*β* (PBL, 12405-1) was added directly to infected cells after spinfection.

For measuring p.f.u., supernatants from infected BMDMs were aspirated, and each well was washed twice with 500 µl sterile milli-Q-grade water. After adding 1 ml of sterile milli-Q water to each well, macrophages were lysed by repeated pipetting. Serial dilutions of lysates were added to confluent Vero cells in 12 well plates. Plates were spun at 300*g* using an Eppendorf 5810R centrifuge for 5 min at room temperature and incubated at 33°C overnight. At ∼16 h.p.i., media was aspirated and replaced with 2 ml/well of DMEM containing 0.7% agarose and 5% FBS (GemCell). At ∼6 d.p.i., 1 ml of DMEM containing 0.7% agarose, 1% FBS (GemCell), 200 µg/ml amphotericin B (Invitrogen, 15290-018), and 2.5% neutral red (Sigma) was added to each well. Plaques were then counted after 24 h.

Microscopy, LDH, and IFN-I experiments were performed as described^21^.

### Animal experiments

Animal research was conducted under a protocol approved by the University of California, Berkeley Institutional Animal Care and Use Committee (IACUC) in compliance with the Animal Welfare Act and other federal statutes relating to animals and experiments using animals (Welch lab animal use protocol AUP-2016-02-8426). The University of California, Berkeley IACUC is fully accredited by the Association for the Assessment and Accreditation of Laboratory Animal Care International and adheres to the principles of the Guide for the Care and use of Laboratory Animals^56^. Mouse infections were performed in a biosafety level 2 facility. All animals were maintained at the University of California, Berkeley campus, and all infections were performed in accordance with the approved protocols. Mice were between 8 and 20 weeks old at the time of initial infection. Mice were selected for experiments based on their availability, regardless of sex. The sex of mice used for survival after i.d. infection and raw data for mouse experiments is indicated in **Supplemental Table 1**. A statistical analysis was not performed to predetermine sample size prior to initial experiments. Initial sample sizes were based on availability of mice and the capacity to process or measure samples within a given time. After the first experiment, a Power Analysis was conducted to determine subsequent group sizes. All mice were of the C57BL/6J background, except for outbred CD-1 mice. All mice were healthy at the time of infection and were housed in microisolator cages and provided chow, water, and bedding. No mice were administered antibiotics or maintained on water with antibiotics. Experimental groups were littermates of the same sex that were randomly assigned to experimental groups. For experiments with mice deficient in *Ifnar* and *Ifngr*, mice were immediately euthanized if they exhibited severe degree of infection, as defined by a core body temperature dropping below 90° F or lethargy that prevented normal movement.

### Mouse genotyping

*Tlr4*^*-/-* 57^, *Ifnar*^*-/-* 58^, *Ifngr*^*-/-* 59^, *Ifnar*^*-/-*^*Ifngr*^*-/-*^ and WT C57BL/6J mice were previously described and originally obtained from Jackson Laboratories. CD-1 mice were obtained from Charles River. For genotyping, ear clips were boiled for 15 min in 60 µl of 25 mM NaOH, quenched with 10 µl Tris-HCl pH 5.5, and 2 µl of lysate was used for PCR using SapphireAMP (Takara, RR350) and gene-specific primers. Primers used were: *Ifnar* forward (F): CAACATACTACAACGACCAAGTGTG; *Ifnar* WT reverse (R): AACAAACCCCCAAACCCCAG; *Ifnar*^*-/-*^ R: ATCTGGACGAAGAGCATCAGG; *Ifngr* (F): CTCGTGCTTTACGGTATCGC; *Ifngr* (R): TCGCTTTCCAGCTGATGTACT; WT *Tlr4* (F): CACCTGATACTTAATGCTGGCTGTAAAAAG; WT *Tlr4* (R): GGTTTAGGCCCCAGAGTTTTGTTCTTCTCA; *Tlr4*^*-/-*^ (F): TGTTGCCCTTCAGTCACAGAGACTCTG; and *Tlr4*^*-/-*^ (R): TGTTGGGTCGTTTGTTCGGATCCGTCG.

### Mouse infections

For mouse infections, *R. parkeri* was prepared by diluting 30%-prep bacteria into cold sterile PBS on ice. Bacterial suspensions were kept on ice during injections. For i.d. infections, mice were anaesthetized with 2.5% isoflurane via inhalation. The right flank of each mouse was shaved with a hair trimmer (Braintree CLP-41590), wiped with 70% ethanol, and 50 µl of bacterial suspension in PBS was injected intradermally using a 30.5-gauge needle. Mice were monitored for ∼3 min until they were fully awake. No adverse effects were recorded from anesthesia. For i.v. infections, mice were exposed to a heat lamp while in their cages for approximately 5 min and then each mouse was moved to a mouse restrainer (Braintree, TB-150 STD). The tail was sterilized with 70% ethanol, and 200 µl of bacterial suspension in sterile PBS was injected using 30.5-gauge needles into the lateral tail vein. Body temperatures were monitored using a rodent rectal thermometer (BrainTree Scientific, RET-3).

For fluorescent dextran experiments, mice were intravenously injected with 150 µl of 10 kDa dextran conjugated with Alexa Fluor 680 (D34680; Thermo Fisher Scientific) at a concentration of 1 mg/ml in sterile PBS^29^. As a negative control, mice with no *R. parkeri* infection were injected with fluorescent dextran. As an additional negative control, uninfected mice were injected intravenously with PBS instead of fluorescent dextran. At 2 h post-injection, mice were euthanized with CO_2_ and cervical dislocation, doused with 70% ethanol, and skin surrounding the injection site (approximately 2 cm in each direction) was removed. Connective tissue between the skin and peritoneum was removed, and skin was placed hair-side-up on a 15 cm Petri dish. Skin was imaged with an LI-COR Odyssey CLx (LI-COR Biosciences), and fluorescence was quantified using ImageStudioLite v5.2.5. The skin from mice with no injected fluorescent dextran was used as the background measurement. Skin from mice injected with fluorescent dextran but no *R. parkeri* was normalized to an arbitrary number (100), and *R. parkeri-* infected samples were normalized to this value (*R. parkeri-*infected / uninfected X 100). The number of pixels at the injection site area was maintained across experiments (7,800 for small area and 80,000 for the large area).

All mice in this study were monitored daily for clinical signs of disease throughout the course of infection, such as hunched posture, lethargy, scruffed fur, paralysis, facial edema, and lesions on the skin of the flank and tail. If any such manifestations were observed, mice were monitored for changes in body weight and temperature. If a mouse displayed severe signs of infection, as defined by a reduction in body temperature below 90°F or an inability to move normally, the animal was immediately and humanely euthanized using CO_2_ followed by cervical dislocation, according to IACUC-approved procedures. Pictures of skin and tail lesions were obtained with permission from the Animal Care and Use Committee Chair and the Office of Laboratory and Animal Care. Pictures were captured with an Apple iPhone 8, software v13.3.1.

For harvesting spleens and livers, mice were euthanized at the indicated pre-determined times and doused with ethanol. Mouse organs were extracted and deposited into 50 ml conical tubes containing 4 ml sterile cold PBS for the spleen and 8 ml PBS for the liver. Organs were kept on ice and were homogenized for ∼10 s using an immersion homogenizer (Fisher, Polytron PT 2500E) at ∼22,000 r.p.m. Organ homogenates were spun at 290*g* for 5 min to pellet the cell debris (Eppendorf 5810R centrifuge). 20 µl of organ homogenates were then serial diluted into 12-well plates containing confluent Vero cells. The plates were then spun at 260*g* for 5 min at room temperature (Eppendorf 5810R centrifuge) and incubated at 33°C. To reduce the possibility of contamination, organ homogenates were plated in duplicate and the second replicate was treated with 50 µg/ml carbenicillin (Sigma) and 200 µg/ml amphotericin B (Gibco). The next day, at approximately 16 h.p.i., the cells were gently washed by replacing the existing media with 1 ml DMEM containing 2% FBS (GemCell). The media were then aspirated and replaced with 2 ml/well of DMEM containing 0.7% agarose, 5% FBS, and 200 µg/ml amphotericin B. When plaques were visible at 6 d.p.i., 1 ml of DMEM containing 0.7% agarose, 1% FBS, and 2.5% neutral red (Sigma) was added to each well, and plaques were counted at 24 h.p.i.

## Statistical analysis

Statistical parameters and significance are reported in the figure legends. For comparing two sets of data, a two-tailed Student’s T test was performed. For comparing two sets of *in vivo* p.f.u. data, Mann-Whitney *U* tests were used. For comparing two survival curves, log-rank (Mantel-Cox) tests were used. For comparing curves of two samples (mouse health, weight, and temperature), two-way ANOVAs were used. For two-way ANOVAs, if a mouse was euthanized prior to the statistical endpoint, the final value that was recorded for the mouse was repeated until the statistical endpoint. For two-way ANOVAs, if a measurement was not recorded for a timepoint, the difference between values at adjacent time points was used. Data were determined to be statistically significant when *P*<0.05. Asterisks denote statistical significance as: \**P*<0.05; \*\**P*<0.01; \*\*\**P*<0.001; \*\*\*\**P*<0.0001, compared to indicated controls. Error bars indicate standard deviation (SD) for *in vitro* experiments and standard error of the mean (SEM) for *in vivo* experiments. All other graphical representations are described in the figure legends. Statistical analyses were performed using GraphPad PRISM v7.0.

## Data availability

WT and *ompB* mutant *R. parkeri* were authenticated by whole genome sequencing and are available in the NCBI Trace and Short-Read Archive; Sequence Read Archive (SRA), accession numbers: SRX4401164 (WT) and SRX4401167 (*ompB*::Tn^STOP^). Raw Data for figures in the main text are available in **Supplemental Table 1**.

## Author contributions

T.P.B. performed and analyzed experiments. C.J.T., P.E., D.R.G., and D.A.E. contributed to performing experiments and provided reagents. T.P.B. wrote the original draft of this manuscript with guidance from M.D.W. Critical reading and edits of the manuscript were provided by C.J.T., P.E., and M.D.W. Supervision was provided by T.P.B. and M.D.W.

## Acknowledgements

We thank Neil Fisher for editing this manuscript. P.E. was supported by postdoctoral fellowships from the Sweden-America Foundation. M.D.W. was supported by NIH/NIAID grants R01AI109044 and R21AI138550. D.R.G., D.A.E., and E.H. were partially supported by NIH/NIAID grant R01 AI24493 (E.H.).

## Supplemental Figures

**Figure S1:**
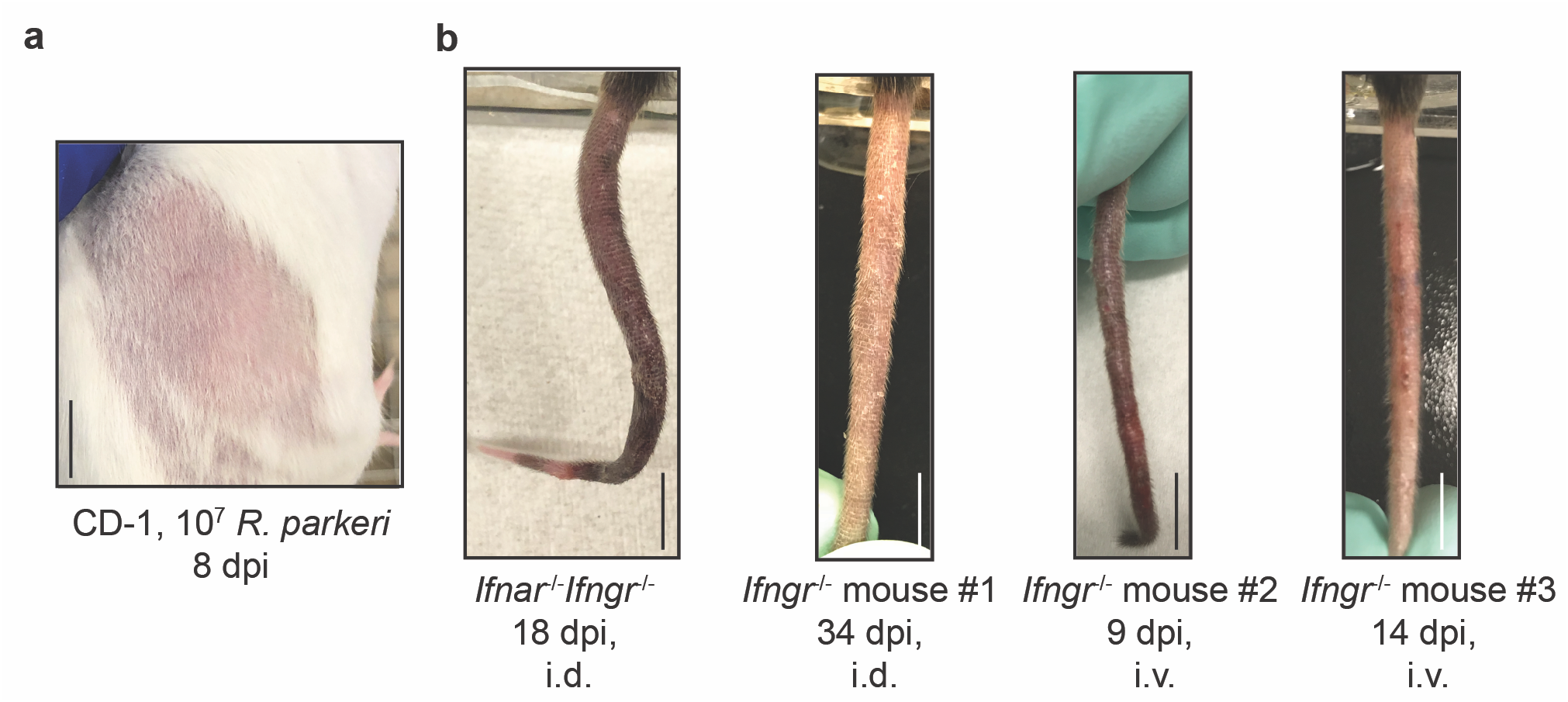
*Ifnar*^-/-^*Ifngr*^-/-^ mice develop disseminated disease upon intradermal *R. parkeri* infection. Representative image of the right flank of CD-1 mice intradermally infected with 10^7^ *R. parkeri*. Scale bar, 1 cm. Data are representative from two independent experiments. Representative images of tails of *Ifnar*^*-/-*^*Ifngr*^*-/-*^ and *Ifngr*^*-/-*^ mice, infected via the i.v. or i.d. route (as indicated), with 10^7^ WT *R. parkeri*. Some *Ifnar*^*-/-*^*Ifngr*^*-/-*^ and *Ifngr*^*-/-*^ mice had no gross pathological manifestations in the tail, whereas some mice exhibited inflamed, necrotic tails at various times post infection. Scale bar, 1 cm. Data are representative from three independent experiments.

**Figure S2:**
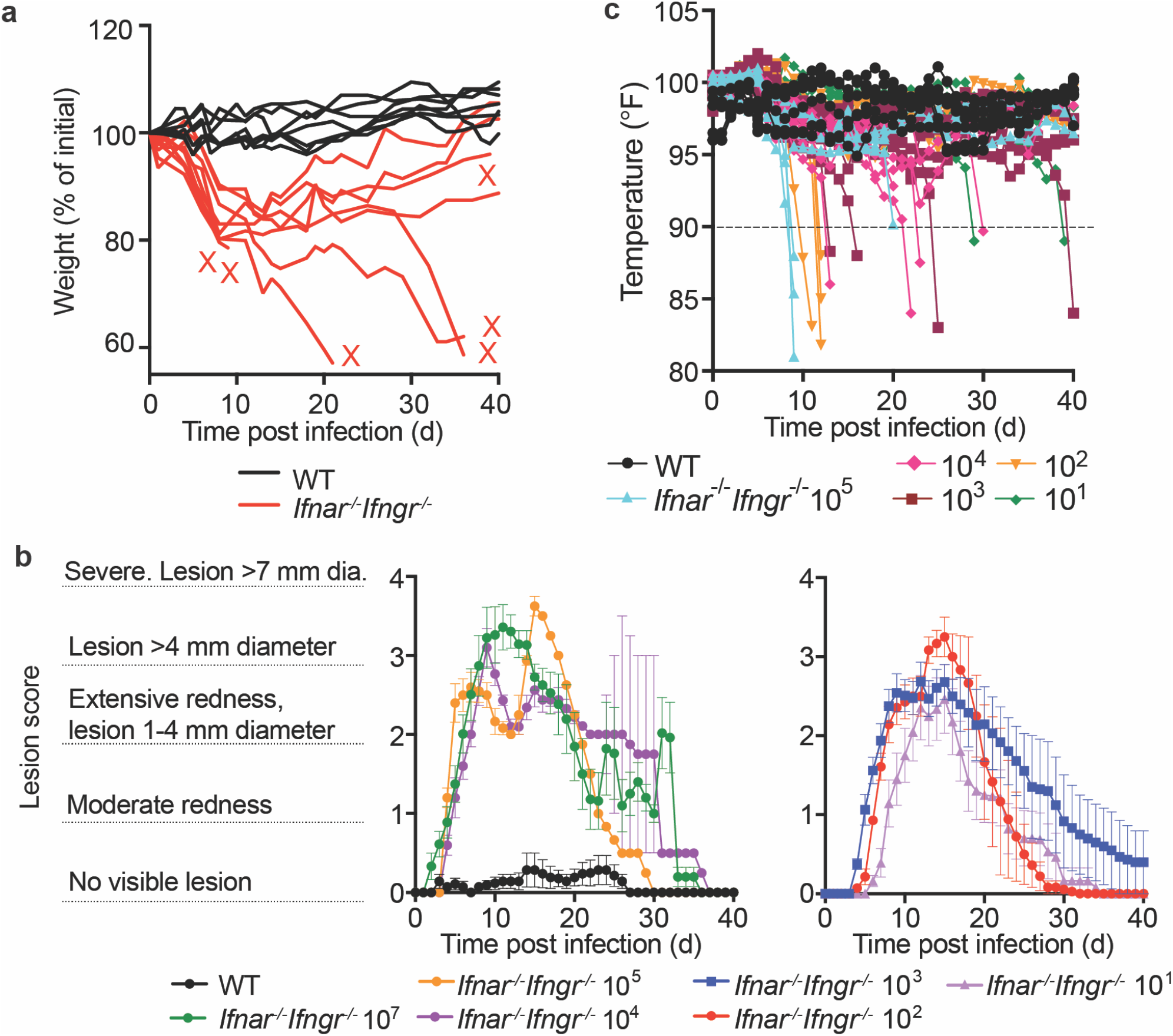
*Ifnar*^*-/-*^ or *Ifngr*^-/-^ mice develop limited disease upon intradermal infection, and *Ifnar*^-/-^ *Ifngr*^-/-^ develop lesions of dose-dependent severity. **a**) Weight changes over time in mice intradermally infected with 10^7^ WT *R. parkeri*. Data are the combination of two independent experiments for WT and three for *Ifnar*^-/-^*Ifngr*^-/-^; *n*=7 (WT) and *n*=9 (*Ifnar*^-/-^*Ifngr*^-/-^) individual mice. Each line is an individual mouse. **b**) Gross pathological analysis of the skin infection site after i.d. infection. *Ifnar*^*-/-*^*Ifngr*^*-/-*^ mice were infected with the indicated number of *R. parkeri* and monitored over time. Data are the combination of three independent experiments for the 10^7^ dose and two independent experiments for all other doses. *n=*7 (WT), *n=*9 (10^7^), *n=*5 (10^5^), *n=*5 (10^4^), *n=*8 (10^3^), *n=*7 (10^2^), and *n=*7 (10^1^) individual mice. Data are the same as in **Fig. 2c** but are extended to 40 d.p.i. Dara are represented as means and error bars indicate SEM. **c**) Temperature changes over time in mice infected i.d. with the indicated amounts of WT *R. parkeri*. Data are the combination of two independent experiments; *n*=7 (WT), *n=*7 (10^5^), *n=*7 (10^4^), *n=*8 (10^3^), *n=*7 (10^2^), and *n=*7 (10^1^) individual mice. Each bar represents an individual mouse. Mice were euthanized if their body temperature fell below 90° F, as indicated by the dotted line.

**Figure S3:**
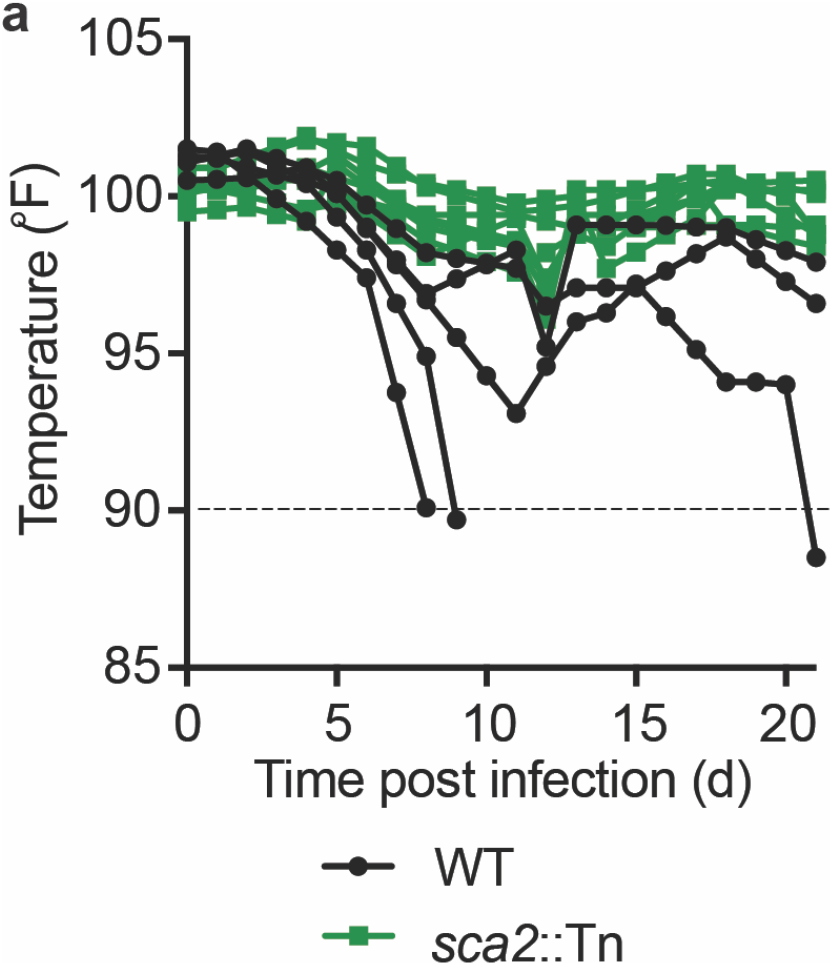
Intradermal infection of *Ifnar*^-/-^*Ifngr*^-/-^ mice with *sca2*::Tn *R. parkeri* causes less severe temperature loss as compared to WT bacteria. **a)** Temperature changes over time in mice infected i.d. with 10^7^ *R. parkeri*. Data are the combination of two independent experiments; *n*=5 (WT), *n=*8 (*sca2*::Tn) individual mice. Each line represents an individual mouse. Mice were euthanized if their body temperature fell below 90° F, as indicated by the dotted line.

**Figure S4:**
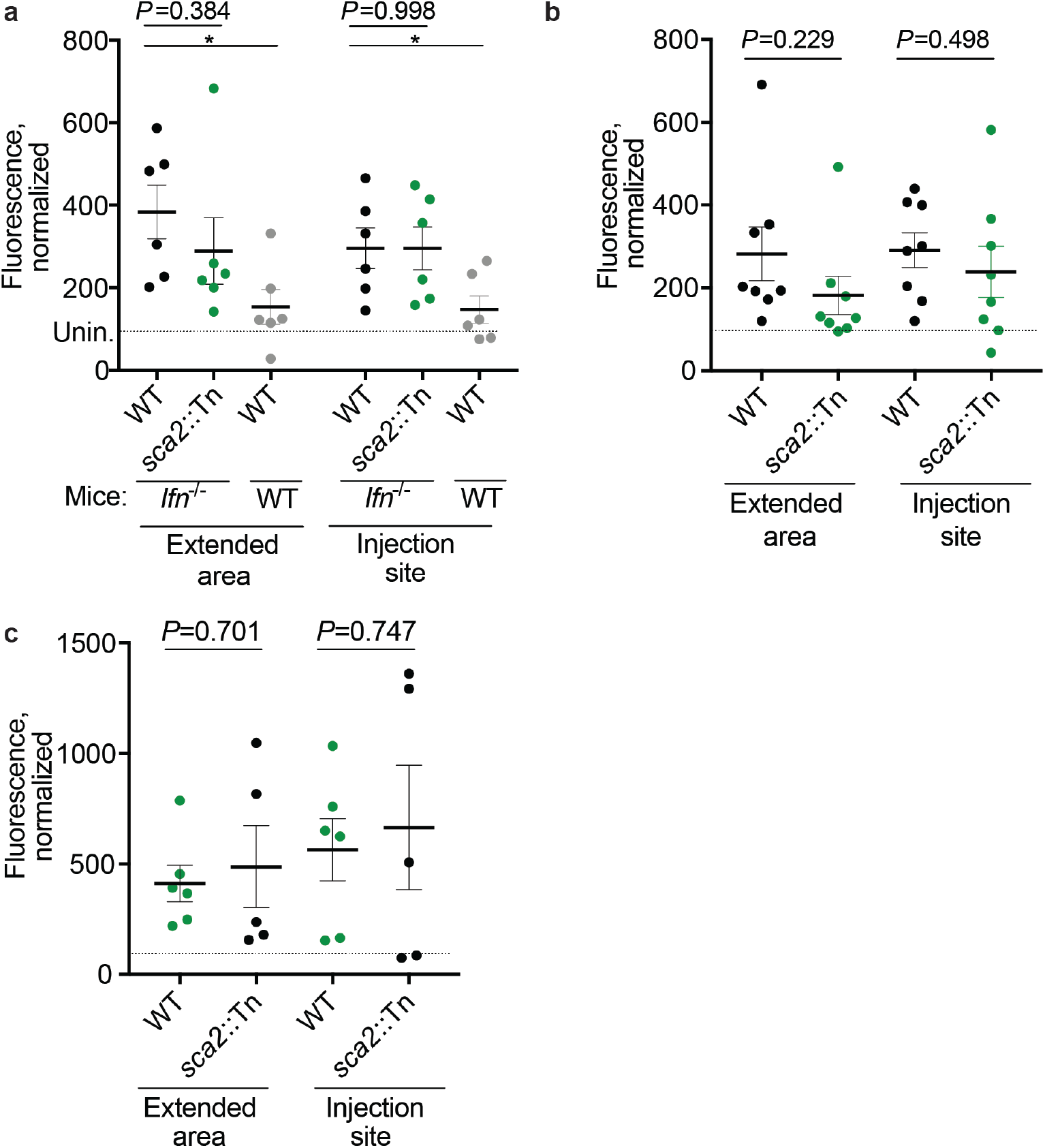
WT and *sca2*::Tn *R. parkeri* elicit similar amounts of vascular damage in skin upon i.d. infection of *Ifnar*^*-/-*^*Ifngr*^*-/-*^ mice. **a**) Quantification of fluorescence in mouse skin after i.d. infection. Mice were infected i.d. with 10^7^ *R. parkeri* and fluorescent dextran was intravenously delivered at 5 d.p.i. Skin was harvested 2 h after delivery of dextran and analyzed with a fluorescence imager. *n*=6 (WT *R. parkeri*) and *n*=6 (*sca2*::Tn *R. parkeri*) individual *Ifnar*^*-/-*^*Ifngr*^*-/-*^ mice; *n*=6 (WT *R. parkeri*) individual WT mice. Data in the ‘extended area’ are the same as those reported in **Fig. 3e. b**) Quantification of fluorescence in mouse skin after i.d. infection. Mice were infected i.d. with 10^6^ *R. parkeri*, and fluorescent dextran was intravenously delivered at 5 d.p.i. Skin was harvested 2 h after delivery of dextran and analyzed with a fluorescence imager. *n*=8 (WT *R. parkeri*) and *n*=8 (*sca2*::Tn *R. parkeri*) individual *Ifnar*^*-/-*^*Ifngr*^*-/-*^ mice. **c**) Quantification of fluorescence in mouse skin after i.d. infection. Mice were infected i.d. with 10^5^ *R. parkeri* and fluorescent dextran was intravenously delivered at 5 d.p.i. Skin was harvested 2 h after delivery of dextran and analyzed with a fluorescence imager. *n*=6 (WT *R. parkeri*) and *n*=5 (*sca2*::Tn *R. parkeri*) individual *Ifnar*^*-/-*^*Ifngr*^*-/-*^ mice. For each experiment, the average of uninfected samples was normalized to 100, and each sample was divided by the average for uninfected mice and multiplied by 100; the dotted horizontal line indicates 100 arbitrary units, corresponding to uninfected (unin.) mice. Representative sizes for the larger ‘extended areas’ of skin and the smaller ‘injection site’ areas of skin are indicated in **Fig. 3d**. Data are each the combination of two independent experiments. Solid horizontal bars indicate means; error bars indicate SEM. For statistical analyses, a two-tailed Student’s T test was used to compare the indicated samples.

**Figure S5:**
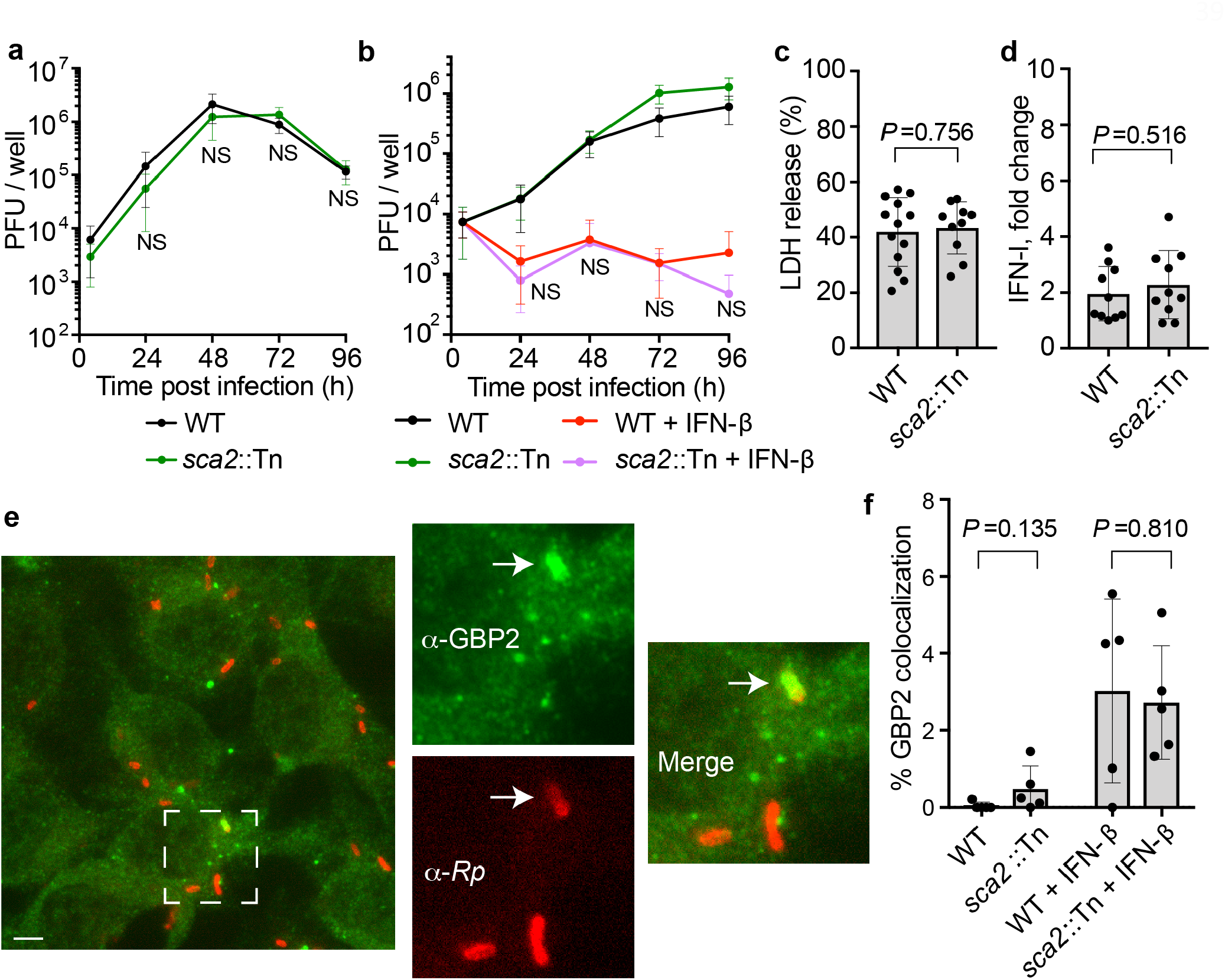
Sca2 does not significantly enhance *R. parkeri* avoidance of antibacterial innate immune responses *in vitro*. **a**) *R. parkeri* abundance in HMEC-1s, multiplicity of infection (MOI) of 0.2. Data are the combination of three independent experiments, each with two biological replicates. For statistics, a two-tailed Student’s T test was used to compare WT to *sca2*::Tn at 48, 72, and 96 h.p.i. No statistically significant differences were observed at any time. **b**) *R. parkeri* abundance in BMDMs, MOI of 1. Data are the combination of three independent experiments, each with two biological replicates. Data were normalized by multiplying fold difference between WT and *Sca2*::Tn at 4 h.p.i. to *Sca2*::Tn and *Sca2*::Tn + IFN-I data at all time points. **c**) Host cell death upon *R. parkeri* infection of BMDMs, as measured by lactate dehydrogenase (LDH) release assay, MOI of 1. From left to right, *n*=6 and 3 biological replicates and are the combination of two independent experiments. **d**) IFN-I abundance in supernatants of infected BMDMs (24 h.p.i.; MOI of 1), measured using a luciferase reporter assay. The data show the fold change over uninfected cells. *n*=7 and 7 biological replicates and are the combination of two independent experiments. **e**) A representative image using x100 confocal immunofluorescence microscopy of WT BMDMs infected with *sca2*::Tn *R. parkeri* in the presence of 100 U recombinant IFN-*β* (3 h.p.i.; MOI of 1). Green, *α*-GBP2; red, *α*-*Rickettsia* (*Rp*). The dotted square indicates the image that is expanded in the other images, separated into two individual and one merged channel. Scale bars, 2.5 µm. White arrows indicate a bacterium that colocalizes with GBP2. Data are representative of three independent experiments. **f**) Quantification of GBP2 colocalization with *R. parkeri* in BMDMs at 24 h.p.i. Each data point is an average of at least five separate images totaling >150 bacteria. Data are the combination of three independent experiments. Statistical analyses used a two-tailed Student’s T test. NS, not significant. Data in **a**,**b** are means; bars in **c, d**, and **f** are means; error bars indicate SD.

